# Legacy of warming and microbial treatments shape root exudates and rhizosphere fungal communities of a tropical tree

**DOI:** 10.1101/2025.08.11.669667

**Authors:** Joel Masanga, Parker Bartz, Gabriela Hernandes Villani, Iana F. Grullón-Penkova, Benedicte Bachelot, Jesse R. Lasky

## Abstract

- Plant metabolites play a pivotal role in shaping rhizosphere microbial communities, yet how plant and microbial functions respond to environmental change remains poorly understood. We combined ecometabolomics and fungal community profiling to investigate how multiple abiotic and biotic treatments influence root exudate chemistry and fungal community assemblage in a tropical tree.
- *Guarea guidonia* seedlings were grown in sterilized soil primed with inoculum from long-term experimentally warmed or ambient plots in Puerto Rico. We manipulated soil microbes, soil moisture, and plant density and tested metabolite-fungi-plant trait associations.
- Metabolite diversity was significantly influenced by soil microbial legacy and moisture, while antimicrobial treatments altered metabolite composition without affecting diversity. Metabolite clusters exhibited distinct treatment-specific patterns. Fungal alpha diversity increased under low moisture, community composition shifted with antimicrobial treatments, and certain families showed strong treatment-specific responses. Although fungal diversity was not correlated with metabolite diversity, fungal community structure was significantly associated with metabolite composition. We also found global and pathway-specific correlations between fungal and metabolite distances, and weak but significant associations between metabolite/fungal composition and seedling traits.
- These results underscore how environmental conditions shape belowground interactions, highlighting metabolite–fungal associations as potential early indicators of plant response to disturbance.

## Introduction

Climate change has intensified disturbance events in the tropics (Emanuel, 2005; Walsh & Ryan, 2000), with major consequences for forest structure, composition, and ecosystem function (Andrewin et al., 2015; Webster et al., 2005; Lugo, 2008). Ecosystem recovery from related disturbances likely hinges on microbial feedbacks, with mutualists enhancing plant resilience and pathogens compounding stress effects (Brooker et al., 2021; Wright et al., 2017). Past evidence from experimental warming and drought studies in tropical forests suggest that such disturbances may for instance reduce pathogen pressure while favoring beneficial microbes (Bachelot et al., 2020), although the biochemical mechanisms mediating these shifts remain unclear. Research on ecological disturbance has largely centered on temperate systems, aboveground traits and single stressors, which not only leaves belowground dynamics underexplored (Melillo et al., 2017; Schindlbacher et al., 2011) but also overlooks the potentially non-additive and complex effects of interacting stressors (Eaton et al., 2020; Robertson & Platt, 2001; Vargas et al., 2010). The Stress Gradient Hypothesis (Bertness & Callaway, 1994) posits that increasing stress favors facilitative over competitive interactions. In this context, Caribbean forests, which are chronically disturbed by storms and warming, offer a valuable system to study how multiple stressors shape belowground ecological dynamics.

Plant-microbe interactions are central to nutrient cycling, seedling establishment, and plant– soil feedbacks (Van Der Heijden et al., 2008; Wagg et al., 2014), yet this key interaction is rarely considered in studies that explore how ongoing changes in climate will shape the future of plant communities. In addition, much of our current understanding of plant-microbe interactions under changing environments comes from crop systems and single-stressor experiments, leaving important gaps in how plants and microbes jointly respond to complex interacting disturbances under natural environments (Trivedi et al., 2022). Plant–microbe–soil interactions heavily rely on root exudates, which contain a repertoire of primary and secondary metabolites that influence rhizosphere microbial composition (Hu et al., 2018) and nutrient cycling (Majumdar et al., 2023; Müller & Junker, 2022; Pang et al., 2021). Further, root exudation and chemistry have been shown to dynamically respond to abiotic and biotic stimuli (Gidman et al., 2003; Gidman et al., 2005; Jones et al., 2011; Lake et al., 2009; Widarto et al., 2006). The interaction between plants, their chemical profiles and rhizosphere microbiome can generate plant–soil feedbacks that influence performance and community dynamics of future communities (Bever, 1994; Gfeller et al., 2023; Hannula et al., 2021; Van der Putten et al., 2013). This might be by altering the soil biotic environment, either positively through recruitment of mutualists such as arbuscular mycorrhizal fungi (AMF) (Bennett et al., 2012; Berendsen et al., 2012; Tamburini et al., 2020), or negatively via pathogen accumulation and resource depletion (Maron et al., 2016; Thakur et al., 2021; Yang et al., 2015). While these interactions have been widely explored in agroecosystems and invasion ecology (Engelkes et al., 2008; Majumdar et al., 2023; van der Putten et al., 2016), their role in mediating tropical forest responses to environmental stress remains underexplored.

Here, we investigated how altered biotic and abiotic conditions influence root exudate chemistry, rhizosphere fungal communities, and seedling traits in the tropical tree species *G. guidonia*. We hypothesized that: (H1) Variation in biotic (microbial alteration) and abiotic (warming, soil moisture and plant density) conditions leads to significant shifts in root exudate metabolic profiles; (H2) These conditions influence rhizosphere fungal community diversity, composition, and abundance; (H3) Differences in root exudate chemistry are associated with corresponding changes in fungal community structure; and (H4) Variation in seedling performance is linked to treatment-driven changes in both root exudate profiles and fungal community composition (Fig. 1a). To test these hypotheses, we employed untargeted metabolomics and amplicon-based fungal community profiling to detect metabolites in root exudates and identify rhizosphere fungal taxa, respectively. We grew *G. guidonia* seedlings in sterilized soil primed with inoculum from either experimentally warmed (+4⍰) or ambient plots at the Tropical Response to Altered Climate Experiment (TRACE) in Puerto Rico, a long-term field warming experiment (Kimball et al., 2018) that has spanned multiple climatic events including major droughts and hurricanes (Alonso-Rodriquez et al., 2022). This design enabled us to assess how microbial communities that resulted from exposure to long-term warming (as a proxy for climate-driven plant-soil-feedback) affect plant metabolite exudation and rhizosphere assembly. We manipulated soil and foliar microbes (via fungicides and heat sterilization), soil moisture (reflecting drought projections described by Bhardwaj et al., 2018), and plant density to test for context-dependent responses. We then evaluated associations between metabolite composition, fungal communities, and seedling traits, with a view that such links reflect early-stage feedbacks relevant to tropical forest resilience and recovery under climate shifts (Jiang et al., 2024; Noecker et al., 2019).

**Fig. 1.**
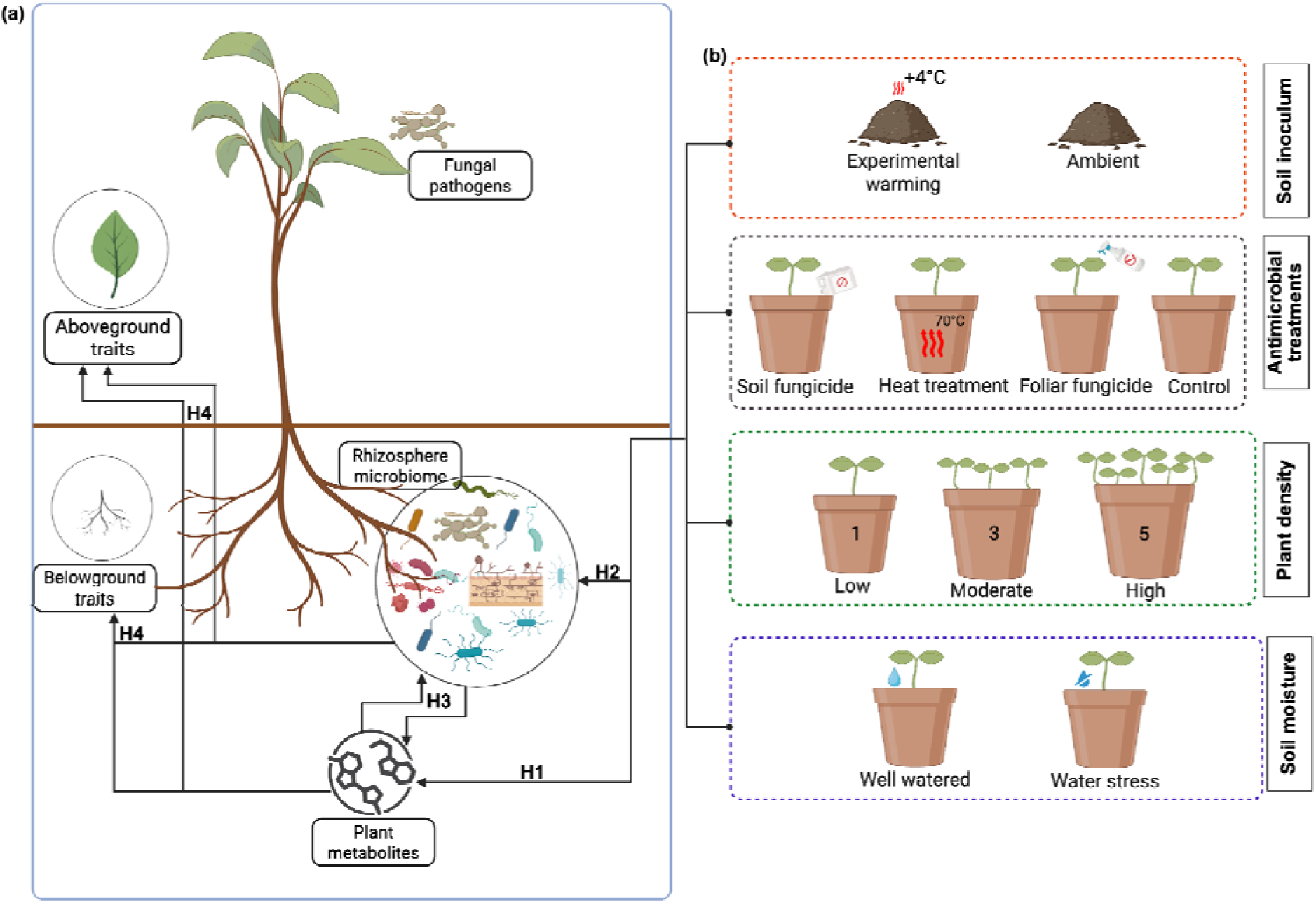
Root exudate metabolomes of *G. guidonia* are shaped by warming-induced microbial legacy and antimicrobial treatments. (a) Metabolite diversity (Functional Hill numbers) acros antimicrobial treatments (x-axis), colored based on microbial legacy conditions (ambient vs. warmed soil). Boxplot whiskers represent 25^th^ and 75^th^ percentiles, while dots indicate biological replicates. (b) NMDS ordination of root exudate metabolite composition based on Bray–Curtis distances of normalized peak intensities. (c) Linear discriminant analysis (LDA) highlighting separation of metabolomic profiles across treatments. (d) Volcano plots of differentially abundant metabolites. Dotted lines indicate raw P-value threshold (P=0.05). Points are colored by FDR-adjusted significance: red = enriched, blue = depleted, grey = not significant.

## Material and Methods

### Experimental treatments, plant material and data collection

This study was conducted under shade house conditions, between January and July 2023 at the USDA Forest Service Sabana Field Research Station in the Luquillo Experimental Forest, northeastern Puerto Rico (18°18⍰N, 65°50⍰W). Forest soil was collected from the neighboring forest, a wet subtropical forest with clay-rich, acidic inceptisols. The soil was sieved, sterilized at 70⍰°C for 48⍰hours, and 250⍰g filled into 4-liter plastic pots. Half of the pots (n = 120) were primed with 5⍰g of soil inoculum collected from either experimentally warmed or ambient control plots at the nearby TRACE field warming experiment. Experimental warming plots have been maintained at +4⍰°C above ambient temperature since 2016, with a one-year pause in warming following Hurricanes Irma and Maria in September 2017 (Reed et al., 2020). These inocula, intended to transfer microbial legacies, are hereafter referred to as “warmed soil” and “ambient soil,” respectively. The pots were further split into four equal batches in which the inoculum was subjected to one of the following antimicrobial treatments; soil fungicide (treated with Banrot™ (Prabhakaran & Dann, 2022); foliar fungicide (treated with Abound Flowable™ (Starkey et al., 2013); heat treatment (oven-sterilized at 70⍰ for 48 hours), and control (no fungicide/heat sterilization treatment). These are collectively referred to as antimicrobial treatments (Fig. 1b). Pots were randomly arranged on raised benchtops, inside a shade house constructed using a gardening tarp that guarantees low light (60% shade) and allows air circulation but restricts insect entry.

We studied the shade tolerant tree *G. guidonia*, a dominant late-successional broadleaved evergreen species common in tropical understories (Pennington and Clarkson 2013). The seedling stage was targeted due to its documented sensitivity to soil microbial variation and environmental stress (Elger et al., 2009). To ensure uniformity across plant material, we grew apparently phenotypically uniform seedlings, germinated from surface sterilized (in 5% NaOCl) seeds collected from mother plants in close proximity. Seedlings were transplanted at low (1 plant/pot), moderate (3 plants/pot), and high (5 plants/pot) densities to test for plant–plant interactions. Water availability was manipulated with water-stress pots receiving 60⍰mL distilled water twice a week and well-watered controls receiving 120⍰mL at the same frequency (Fig. 1b). After three months of growth, root exudates were collected by exposing a single lateral root (still attached to the plant) in each pot, gently cleaning it with ethanol-soaked cotton balls, and submerging it in 5⍰mL deionized water for 48⍰hours under darkness provided by foil-wrap. Samples were stored at −80⍰°C until metabolomic profiling. Plants were then phenotyped for aboveground traits (leaf surface area, specific leaf area, and leaf number), prior to harvesting. Roots were scanned using WinRHIZO™ (Regent Instruments Inc., Canada) to assess total root length, root surface area, root diameter, root volume, and number of root tips. Subsamples were flash-frozen at −80⍰°C for rhizosphere fungal community sequencing.

### Liquid Chromatography-Mass Spectrometry

Root exudates were centrifuged (300⍰rpm, 20⍰min) and concentrated using solid-phase extraction (SPE) prior to LC–MS/MS analysis. Samples were passed through C18 columns (HyperSep SPE, Thermo Fisher, USA) preconditioned with acidified methanol (1% formic acid) and HPLC-grade water under vacuum (Resprep manifold, Restek). Metabolites were eluted in 3⍰mL methanol, concentrated in a centrifugal vacuum concentrator (SpeedVac), and reconstituted in 100⍰µL of 60:40 methanol:water containing 0.1⍰µM chlorpropamide (internal standard). LC–MS/MS was performed at the Huck Metabolomics Core Facility, Pennsylvania State University, on a Thermo Vanquish Horizon system (Thermo Fisher Scientific, Waltham, MA, USA) equipped with a Waters BEH C18 column (2.1 × 150⍰mm, 1.7⍰μm) at 55⍰°C. The mobile phase consisted of 0.1% formic acid in water (solvent A) and 0.1% formic acid in acetonitrile (solvent B), with the following gradient: 10% B (0–2⍰min), ramped to 40% B (12⍰min), then to 100% B (15⍰min), held until 21⍰min, returned to 10% B at 22⍰min, and re-equilibrated until 30⍰min. The flow rate was 0.3⍰mL/min. The eluate was introduced into a Thermo Orbitrap Exploris 120 mass spectrometer using heated electrospray ionization. Full scan data (m/z 100–1000) were acquired at 120,000 resolution, followed by ddMS2 scans at 15,000 resolution using normalized collision energies of 15, 30, and 45⍰V. Capillary voltages were 2500⍰V (negative mode) and 3500⍰V (positive mode). Sheath gas was set to 40, auxiliary gas to 6, and sweep gas to 0. The ion transfer tube and vaporizer were set to 325⍰°C and 350⍰°C, respectively. Data was obtained in both positive and negative modes, and analyzed using MetaboAnalyst 6.0 (Pang et al., 2024).

### Fungal community characterization

To assess fungal community structure, we sampled roots from pots with surviving seedlings following exudate collection at the end of the 3 months growth period. Root samples were lysed using a TissueLyser II (Qiagen, Germantown, MD, USA) with ceramic beads, and total DNA extracted using the PowerPlant Pro Kit (Qiagen), following the manufacturer’s protocol with minor modifications. PCR amplification was performed targeting the fungal internal transcribed spacer 2 (ITS2) region using primers ITS4_Nextera and JL0015.8SR_Nextera. PCR cycling conditions were: 95⍰°C for 3⍰min (initial denaturation), followed by 30 cycles of 98⍰°C for 20⍰s, 65.7⍰°C for 15⍰s, and 72⍰°C for 45⍰s, with a final extension at 72⍰°C for 5⍰min (Gohl et al., 2016). Sequencing was performed on the NextSeq P1 XLEAP platform at the University of Minnesota Genomic Center (UMGC). Initial quality control and demultiplexing were also conducted by UMGC. Sequence processing and operational taxonomic units (OTU) clustering were performed on the Pete supercomputer at Oklahoma State University using a modified Mothur pipeline (Schloss et al., 2009). After detecting and removing chimeras, this pipeline generates operational taxonomic units (OTUs) by grouping sequences at 97% identity (Gweon et al., 2015) using the UNITE v9 database (Abarenkov et al., 2024). Taxonomic classification and ecological guild assignment were completed using FUNGuild (Nguyen et al., 2016), enabling annotation of fungal OTUs by functional role.

### Statistical analyses

#### Treatment effects on metabolite diversity and composition

We analyzed exudate metabolite diversity using the ‘chemodiv’ R package (Petrén et al., 2023), which calculates Functional Hill numbers integrating compound richness, relative abundance, and structural dissimilarity (Chiu & Chao, 2014). Compounds were categorized using the function *NPCTable*, and pairwise dissimilarities computed with *compDis*. Diversity metrics were compared across antimicrobial treatments, microbial legacy, plant density, and soil moisture using two-tailed t-tests or one-way ANOVA. To assess treatment effects on metabolite composition, we computed Bray–Curtis dissimilarity matrices and conducted PERMANOVA using *adonis2* function in ‘vegan’. We used Linear Discriminant Analysis (LDA), implemented in the ‘LDA’ package to identify metabolites that best discriminated among treatment groups. Significant contributors (p < 0.05) were ranked based on LDA scores. Differential abundance of individual metabolites was evaluated using the ‘limma’ package, relative to control (untreated inoculum) as the reference group. Metabolites were considered significantly modulated if they had a false discovery rate (FDR) < 0.05. To explore non-linear treatment effects, we employed the package ‘cluster’ (Freire-Zapata et al., 2024) to identify metabolite clusters based on co-occurrence patterns across samples. Only metabolites detected in ≥50% of samples were included. Pairwise dissimilarities were computed based on Manhattan distances using the function *daisy*, while clustering was performed using Partitioning Around Medoids (PAM) using the function *pam*. The optimal number of clusters (k) was determined by silhouette analysis. Consensus values, reflecting sample-level contributions to each cluster, were calculated for use in downstream correlations with fungal diversity and seedling traits.

#### Treatment effects on fungal diversity, composition and differential abundance

To characterize fungal diversity across experimental treatments, we calculated *alpha* diversity indices from OTU tables after rarefaction using the package ‘phyloseq’ (McMurdie & Holmes, 2013). These metrics were compared across treatments using t-tests or ANOVA. Fungal *beta* diversity was estimated via Bray–Curtis dissimilarity and tested using PERMANOVA. Differential abundance analysis was conducted using the package ‘ANCOM-BC’ (Lin & Peddada, 2020), which accounts for compositional bias and sample-level heterogeneity. Taxa were compared relative to controls, and significance was determined at FDR<0.05 based on the Benjamini– Hochberg correction.

#### Metabolite-fungal community associations

We examined metabolite–fungal associations using multiple complementary approaches. (i) Mantel tests, conducted in ‘vegan’ with 999 permutations, were performed between Bray– Curtis dissimilarity matrices of metabolites and fungal communities for all samples that had both datasets (n = 144). This included all detected compounds (including unknowns, n=612) and pathway subsets, namely alkaloids (n=150), amino acids and peptides (n=68), carbohydrates (n=28), fatty acids (n=91), polyketides (n=23), shikimates and phenylpropanoids (n=110), and terpenoids (n=79). (ii) Partial Spearman’s rank correlations, implemented in the ‘ppcor’ v.1.1 package (Kim, 2015) were used to test associations between individual metabolite features and fungal Shannon diversity and OTU abundance. All models were adjusted for microbial legacy, antimicrobial treatment, plant density, and soil moisture. Results were corrected for multiple comparisons using FDR. (iii) Redundancy Analysis (RDA) was conducted using the first 10 metabolite principal components (PCs) to quantify explained variance in fungal community composition. Contributions of individual PCs to RDA axes were assessed via biplot loadings, while outlier fungal taxa associated with metabolite gradients were identified based on Mahalanobis distances from the RDA centroid. These taxa were interpreted as drivers of community-level metabolomic associations. (iv) Spearman’s rank tests, implemented in the ‘Hmisc’ package, were used to correlate metabolite cluster consensus values with fungal Shannon diversity, and results corrected for multiple comparisons using FDR.

#### Metabolite- and fungi-seedling trait associations

To test whether variation in metabolite or fungal composition was linked to seedling traits, we first constructed a Euclidean distance matrix from six morphological traits (root surface area, root diameter, root volume, dry root mass, leaf surface area, and specific leaf area). This matrix was correlated with metabolite and fungal community dissimilarities using Mantel tests. To detect coordinated divergence, we extracted sample pairs in the upper-right quadrant (top 25%) of both trait vs metabolite and trait vs fungal Mantel test results. Treatment combinations within these high-divergence pairs were summarized in contingency tables and visualized as heatmaps to identify dominant treatment contrasts driving divergence. Finally, to detect metabolite–trait associations beyond global metabolomic distance metrics, we correlated metabolite cluster consensus values (from cluster analysis) with individual plant traits using Spearman’s rank tests, followed by FDR correction.

## Results

### Warming-induced microbial legacy and heat sterilization of soil inoculum influence exudate metabolomes

We hypothesized that biotic (microbial alteration) and abiotic (warming, soil moisture and plant density) conditions and/or their combinations influence root exudate metabolite profiles in *G. guidonia* (H1). To test this hypothesis, we conducted untargeted metabolomics on root exudates from 162 individuals and identified 612 putative metabolites (471 in positive and 141 in negative ion modes). NPClassifier assigned the compounds to seven major pathways: alkaloids (150), amino acids and peptides (68), carbohydrates (28), fatty acids (91), polyketides (23), shikimates and phenylpropanoids (110), and terpenoids (79). Notably, 63 features could not be classified. Next, we quantified metabolite diversity using Functional Hill numbers and compared diversity across treatments. Significantly higher metabolite diversity was observed under ambient than warmed soil inoculum (t = 3.71, p = 0.0003), and under high than low soil moisture (t = 2.12, p = 0.036) (Fig. 2a; Fig. S1). In contrast, neither antimicrobial treatments nor plant density significantly affected metabolite diversity (both p > 0.05). Likewise, antimicrobial treatments did not alter the overall distribution of metabolic pathways (Fig. S1). Metabolite composition was significantly influenced by microbial legacy and antimicrobial treatments (p < 0.05 for both), but not by soil moisture or plant density (p > 0.05) (Fig. 2b). Together, these results suggest that warming legacy effects on microbes and soil moisture level primarily influence metabolite richness. In contrast, microbial alteration and plant density have a stronger effect on metabolite composition, shifting the relative abundances of specific compounds rather than the total number present.

**Fig. 2.**
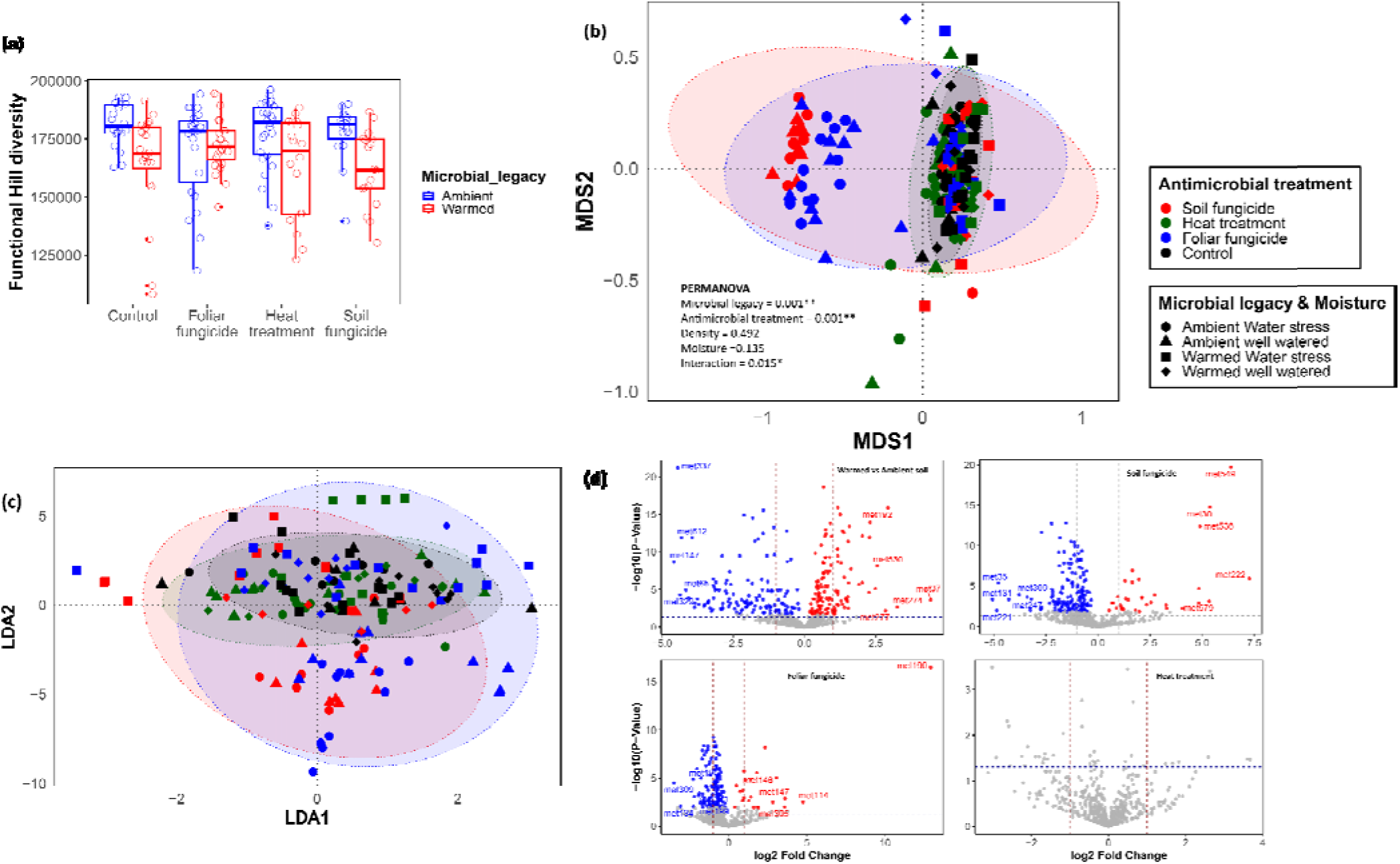
Diversity and composition of rhizosphere fungal communities associated with *G. guidonia*. (a) Shannon diversity of fungal communities across antimicrobial treatments (x-axis) and moisture. H and L denote high and low moisture levels, respectively. (b) Relative abundance of dominant fungal families across antimicrobial treatments. (c) Principal coordinate analysis (PCoA) based on Bray–Curtis dissimilarities illustrating shifts in fungal community composition in response to treatments. (d) Differential abundance of fungal taxa based on ANCOM-BC analysis. Enrichment and depletion patterns are shown relative to the untreated control group.

To further understand how treatments shaped metabolite composition, we performed linear discriminant analysis (LDA), and observed clear separation based on microbial legacy (Fig. 2c). The top 20 metabolites contributing to each LDA axis are shown in Fig. S1, while the full list of differentially discriminant features is provided in Dataset S1. To identify specific compounds driving these patterns, we applied the ‘limma’ package in R for differential enrichment and depletion of metabolites under microbial legacy and antimicrobial treatments, two factors that significantly influenced composition (Fig. 2b). We identified 255 metabolites that differed significantly between warmed and ambient inoculum sources (126 enriched and 129 depleted; FDR < 0.05). Additionally, 183 and 148 metabolites were differentially abundant in soil fungicide and foliar fungicide treatments, respectively, compared to untreated controls (FDR < 0.05). The heat-treated condition yielded 39 metabolites that were nominally significant (P < 0.05) but did not pass FDR correction (FDR > 0.05) (Fig. 12; Fig. S2).

### Metabolite clusters reveal distinct treatment-specific responses

We identified three metabolite clusters based on co-occurring features. The clusters consisted of 88 (Cluster 1), 141 (Cluster 2), and 53 (Cluster 3) metabolites, with all seven annotated metabolite pathways represented across clusters (Fig. S2). Heatmaps revealed clear treatment-specific shifts in metabolite abundance relative to control conditions across the three clusters (Fig. S2). To further evaluate treatment effects, we fitted linear models with interaction terms and found that metabolites in Cluster 1 were significantly influenced by all experimental treatments (p < 0.05) except low plant density (p > 0.05). Cluster 2 metabolites responded significantly to microbial legacy (p = 0.004), antimicrobial treatments (p < 0.001), and moisture levels (p < 0.001). Cluster 3 metabolites showed a similar treatment response to Cluster 1. Linear model results are summarized in Dataset S2-S4. By grouping metabolites based on co-occurrence, we uncovered nuanced environmental effects that were potentially obscured in traditional single-metabolite models.

### Soil moisture and antimicrobial treatments influence rhizosphere fungal diversity and community composition

Amplicon sequencing and rarefaction identified 359 fungal OTUs in the *G. guidonia* rhizosphere, spanning 8 phyla, 18 classes, 59 orders, 130 families, 144 genera, and 200 species. Fungal Shannon diversity was significantly higher under low moisture relative to well-watered conditions (t = 2.33, p = 0.022; Fig. 3a). In contrast, no significant differences in Shannon or other *alpha* diversity metrics were observed across microbial legacy, antimicrobial, or plant density treatments (p > 0.05; Fig. S3–S4). At the family level, antimicrobial treatments induced compositional shifts, with distinct variation in fungal taxa across treatments (Fig. 3b). To assess fungal community responses to experimental treatments, we applied principal coordinate analysis (PCoA) on Bray–Curtis dissimilarity matrices followed by PERMANOVA. Fungal community composition differed significantly across antimicrobial treatments (p = 0.008), but not across soil inoculum source, plant density, or moisture when considered individually (p > 0.05 for all). However, a significant interaction effect (p = 0.005; Fig. 3c) emerged when all treatment variables were considered jointly, suggesting that fungal assemblages are shaped by the combined influence of combined environmental alteration.

**Fig. 3.**
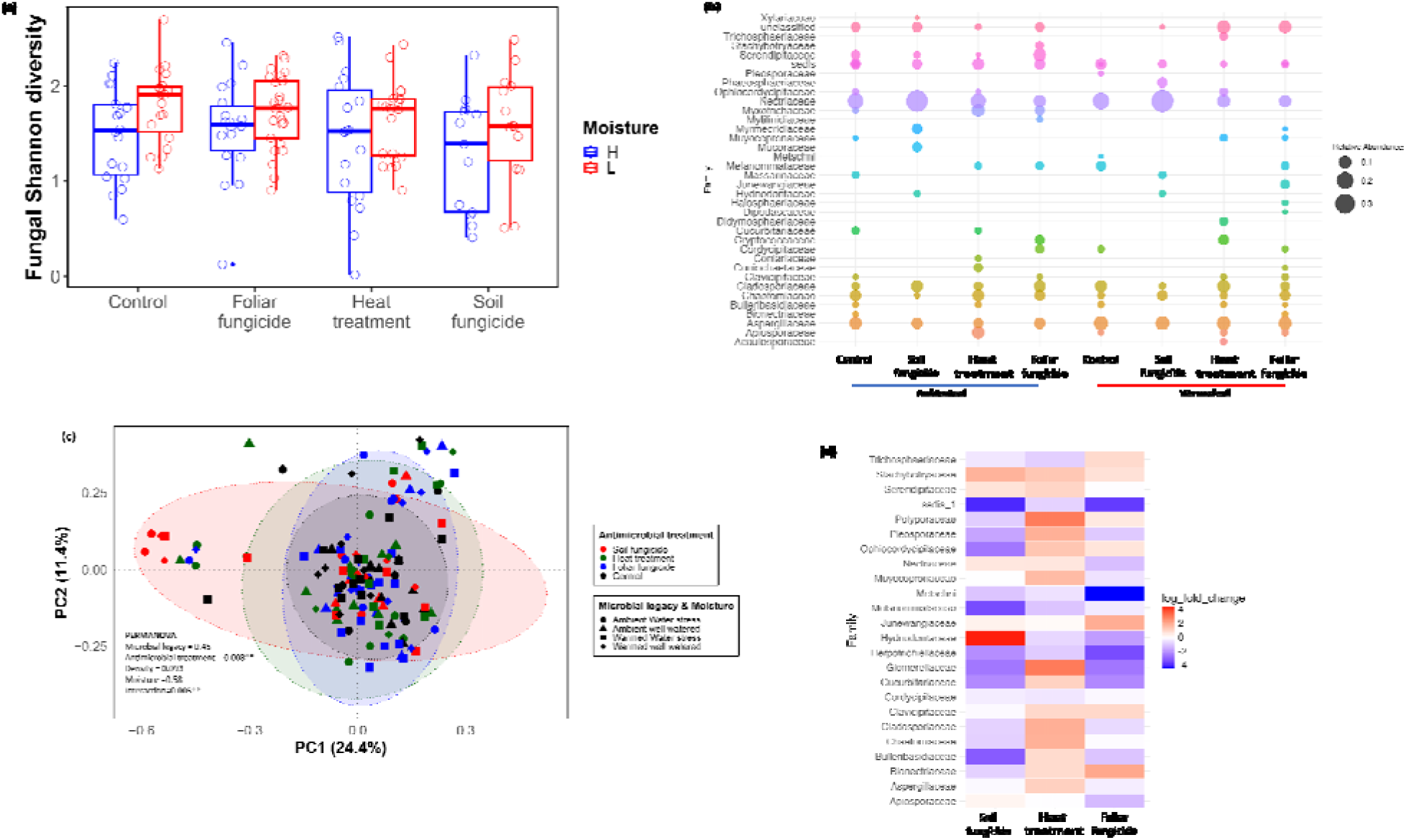
Associations between fungal communities and root exudate metabolites. (a–h) Mantel test results showing correlations between fungal Bray–Curtis dissimilarity with (a) all metabolites, (b–h) individual metabolite classes (x-axis). (i) Partial Spearman’s rank correlation between fungal Shannon diversity and individual metabolites, adjusted for soil inoculum, antimicrobial treatments, soil moisture, and plant density. Red-positive associations; Blue: negative associations. Dotted line indicates p = 0.05 threshold. Top 3 positively and negatively associated metabolites are labeled. *-6,10a-dihydroxy-4-(hydroxymethyl)-4,7,11b-trimethyl-1,2,3,4a,5,6,6a,7,11,11a-decahydronaphtho[2,1-f][1]benzofuran-9-one.

Next, we used ANCOM-BC to examine family-level shifts in fungal differential abundance acros antimicrobial treatments, controlling for microbial legacy, plant density, and soil moisture. Taxa from control pots (no treatment on soil inoculum) served as the baseline for differential abundance analysis. We found that the soil fungicide treatment resulted in elevated abundance of Hydnodontaceae and Stachybotryaceae, while heat sterilization caused significant enrichment of Polyporaceae and Glomerellaceae. The foliar fungicide treatment increased the abundance of Bionectriaceae and Junewangiaceae. Notably, Hydnodontaceae was enriched under soil fungicide but depleted in heat and foliar fungicide treatments, wherea Glomerellaceae showed an opposite trend, enriched under heat treatment but depleted by fungicides. These patterns (Fig. 3d) support our earlier findings that antimicrobial treatments distinctly shape rhizosphere fungal assemblages.

### Metabolite composition is associated with fungal community structure

To assess the relationship between rhizosphere metabolite composition and fungal community structure, we used redundancy analysis (RDA), with the first 10 metabolite principal components (PCs) as explanatory variables. These PCs captured most of the variation in metabolite profiles while reducing dimensionality. The RDA model explained a modest proportion of fungal community variation (R² = 0.0614; Adjusted R² = –0.00008; Fig. S5a), with PC2 and PC10 contributing most to RDA1, and PC1 and PC4 to RDA2. Most PCs were characterized by predominantly positive metabolite loadings, suggesting that metabolite patterns captured by these axes were broadly associated with shifts in fungal composition (Fig. S5b). While PCs reflect underlying metabolite structure, RDA scores reflect taxon responses to those multivariate gradients. References to PC contributions to RDA axes illustrate the strength of association between metabolite-derived variation and community composition, rather than a direct mapping of metabolite loadings onto fungal taxa. Outlier fungal taxa, identified via Mahalanobis distance, included 64 OTUs. These taxa were predominantly from phylum Ascomycota (84%), with Nectriaceae making up over half of the total (Fig. S5c). To complement this multivariate approach, we used Mantel tests to assess the strength of correlation between fungal community distances and metabolite dissimilarity. We found a significant positive correlation between global metabolite composition (all 612 features) and fungal community structure (r = 0.006, p < 0.001; Fig. 4a). Importantly, all seven metabolite classes showed statistically significant associations with fungal distances (p <0.05 for all comparisons; Fig. 4b–h). These results indicate a strong covariation between rhizosphere fungal communities and exuded metabolite profiles, consistent with potential reciprocal influences between plant chemistry and microbial community composition. To explore whether metabolite profiles were associated with fungal diversity rather than just composition, we used partial Spearman’s rank correlation models adjusted for soil inoculum source, antimicrobial treatments, soil moisture, and plant density. We identified several positive and negative metabolite associations with fungal Shannon diversity (Fig. 4i). However, none remained significant after multiple testing (FDR >0.05). We also checked for additional metabolite vs fungal association patterns, focusing on consensus metabolite cluster values, and observed that Clusters 2 and 3 exhibited significant positive correlations with fungal Shannon diversity (p < 0.05), whereas Cluster 1 had a wea negative association (Table S1).

**Fig. 4.**
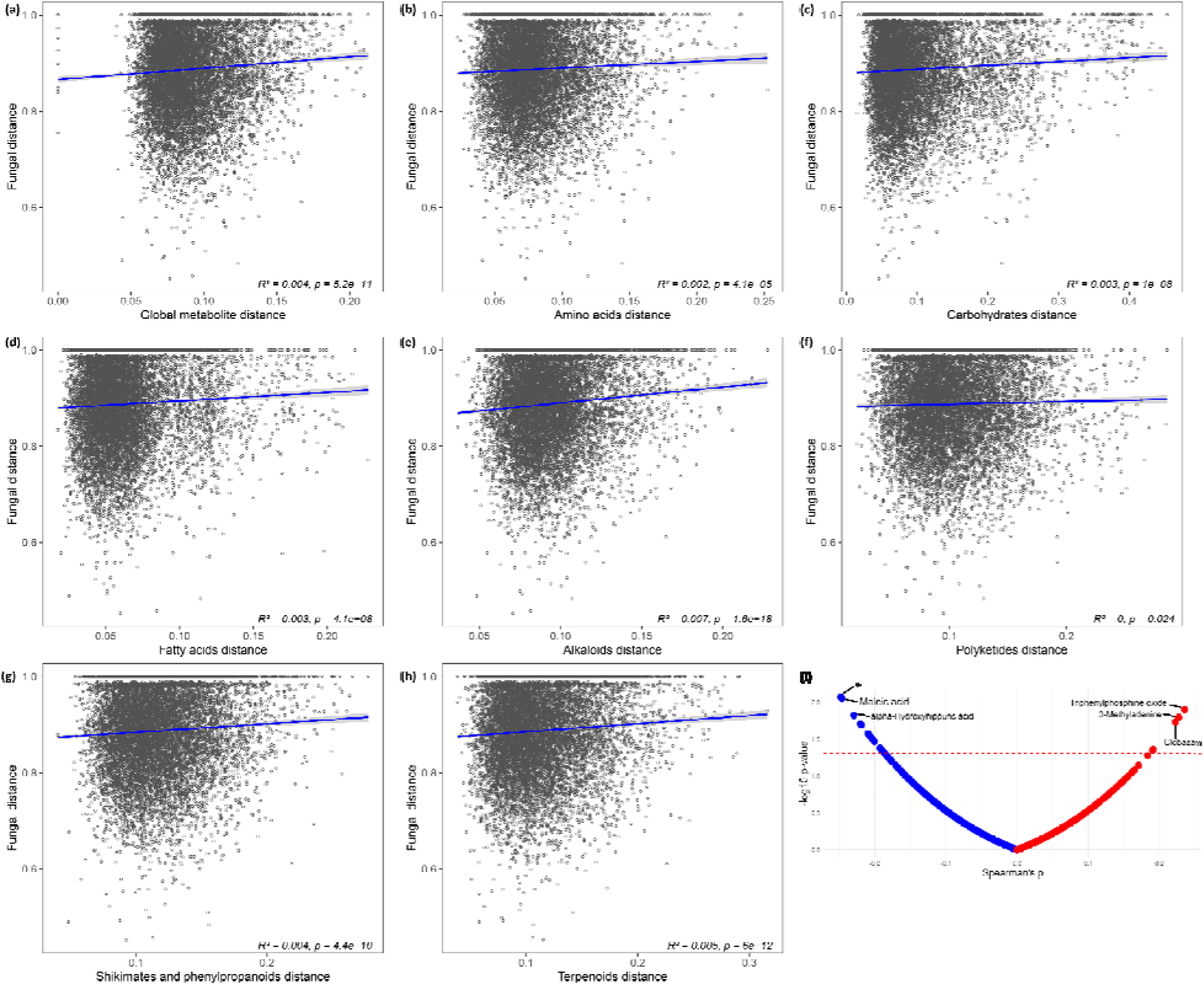
Associations between seedling trait divergence and metabolite and fungal community dissimilarity. (a–b) Mantel test results showing pairwise correlations between seedling trait distances with (a) root exudate metabolite dissimilarity and (b) fungal community dissimilarity. (c–d) Heatmaps summarizing the frequency of treatment pairings contributing to high-divergence sample pairs extracted from the upper 25% quantile (≥75 percentile) for both trait vs metabolite or fungal distances. Warmer colors represent greater co-occurrence of treatment combinations among highly divergent samples, highlighting treatment-driven shifts in associations.

### Association between seedling traits with exudate metabolites and rhizosphere fungal communities

All studied seedling traits, with the exception of dry root mass, did not significantly vary with experimental treatments (Fig. S6-7 and Table S2). To test whether belowground processes were linked to trait expression (H4), we correlated seedling traits with exudate metabolite and fungal community data. Seedling traits showed no significant association with metabolite diversity (Functional Hill index). In contrast, trait dissimilarity was significantly correlated with metabolite composition (Mantel R² = 0.018, p < 0.001) (Fig. 5a), suggesting that compositional shifts mirror morphological variation more strongly than diversity metrics alone. With regards to the metabolite clusters, leaf surface area exhibited the strongest association with consensus cluster values, negatively correlating with Cluster 1 and positively with Clusters 2 and 3 (FDR-adjusted p<0.005), suggesting possible trade-offs in carbon dynamics. Associations with dry root mass and specific leaf area were weaker and less consistent (Table S1). Fungal community composition was significantly correlated with trait distance (Mantel R² = 0.001, p = 0.0031; Fig. 5b), while no association was observed between fungal Shannon diversity and seedling traits (p >0.05).

**Fig. 5.**
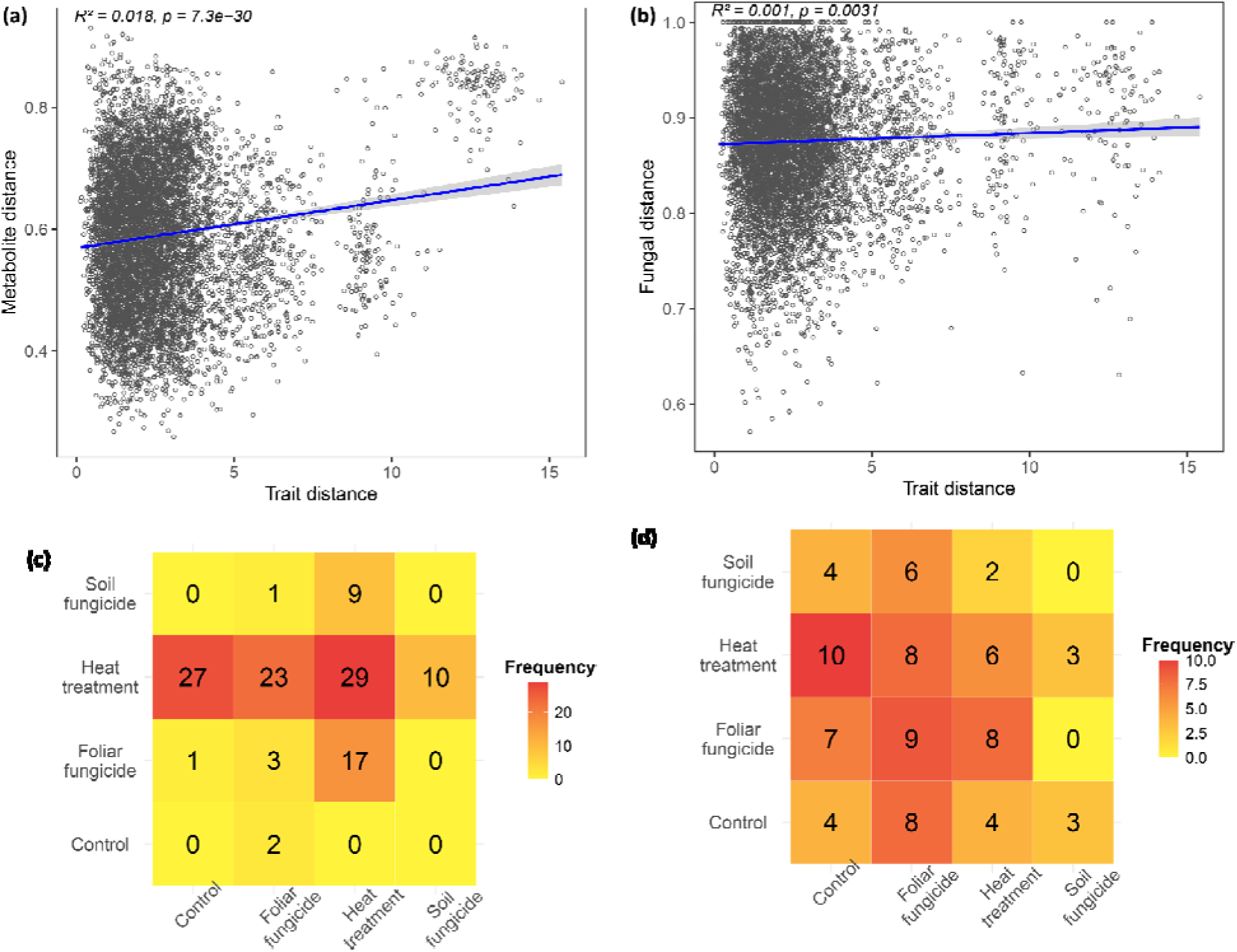
Associations between seedling trait divergence and metabolite and fungal community dissimilarity. (a–b) Mantel test results showing pairwise correlations between seedling trait distances with (a) root exudate metabolite dissimilarity and (b) fungal community dissimilarity. (c–d) Heatmaps summarizing the frequency of treatment pairings contributing to high-divergence sample pairs extracted from the upper 25% quantile (≥75^th^ percentile) for both trait vs metabolite or fungal distances. Warmer colors represent greater co-occurrence of treatment combinations among highly divergent samples, highlighting treatment-driven shifts in associations.

To further assess treatment-driven divergence, we compared high-dissimilarity sample pairs from Mantel plots, focusing on points in the ≥75^th^ percentile in both axes across metabolite/trait and fungal/trait associations. Trait–metabolite divergence was strongest in comparisons involving heat-treated inoculum (particularly when compared to control and foliar fungicide treatments), highlighting a strong legacy effect of microbial depletion or abiotic stress on exudation and plant development. By contrast, soil fungicide treatments showed little trait–metabolite divergence, suggesting that AMF-specific disruption had limited impact. Trait–fungal divergence was more broadly distributed, with mild increases in pairs involving foliar fungicide and heat treatments (Fig. 5c–d). Overall, these results suggest that root exudate metabolite are more tightly linked to seedling trait variation than fungal communities, with treatment history potentially playing a key role in shaping these interactions.

## Discussion

Tropical forests face simultaneous environmental disturbances (Bennett et al., 2021; Lugo, 2008), whose impacts have broad implications for global ecosystem function. While aboveground biodiversity responses are more frequently studied (Malhi et al., 2014), belowground processes-particularly those linking root exudation, microbial assembly, and soil biotic legacies-remain underexplored. Root exudates are key drivers of rhizosphere microbiome composition and can influence plant resilience to biotic and abiotic stresses (Badri & Vivanco, 2009; Canarini et al., 2019), yet how climate-induced shifts in microbial communities feed back into these interactions remains poorly understood.

Here, we examined how warming-induced soil microbial legacy influences metabolite composition in root exudates and associated belowground interactions in a tropical tree species. Using inoculum from experimentally warmed and ambient plots, combined with manipulations of fungal communities, plant density, and soil moisture, we found that exudate metabolite–fungi relationships are shaped by specific treatments rather than strong and broad global correlations. Metabolite diversity was significantly influenced by warming-induced microbial legacy and high moisture, whereas composition shifted with microbial legacy and antimicrobial treatments. Variations in metabolite profiles coincided with shifts in fungal community structure, particularly among families such as Hydnodontaceae, Glomerellaceae, and Nectriaceae. In addition, metabolite classes were significantly correlated with fungal composition. While fungal communities had modest relationships with seedling traits, metabolite profiles exhibited weak but significant associations with trait variation, particularly leaf surface area. Together, our results point to a context-dependent relationship between root exudate metabolites and fungal communities, structured by both abiotic stress and microbial legacy. These findings underscore the complexity of belowground interactions in tropical systems and highlight root exudates as early biochemical signals in plant responses to environmental change.

### Biotic and abiotic treatments alter plant metabolic profiles

Environmental change is a powerful driver of belowground ecological dynamics, shaping both soil microbial communities and the diverse repertoire of plant metabolites released in root exudates in response to abiotic and biotic stressors (Bardgett & Caruso, 2020; Hu et al., 2018; Ma et al., 2022; Pang et al., 2021; Trivedi et al., 2022; Van der Putten et al., 2013; Yang et al., 2025). Our results show that G. guidonia responds to environmental alterations through changes in metabolite exudation and rhizosphere fungal community structure. Specifically, microbial legacy emerged as a key determinant of metabolite composition, supporting the idea that long-term changes in microbial communities can influence the synthesis and secretion of a wide range of plant metabolites, with potential feedback effects on the rhizosphere microbiome (Cao et al., 2020; Wagg et al., 2014). It is important to note that some of the exuded compounds may originate from microbial metabolism or transformation of plant-derived compounds, especially under fluctuating environmental conditions. These shifts likely reflect plasticity in plant chemical investment, potentially mediated by altered nutrient cycling or microbial signaling that regulate secondary metabolism (Badri & Vivanco, 2009; Canarini et al., 2019). Microbe-altering treatments further modified exudate composition, possibly through feedbacks that reconfigure carbon allocation to defense or nutrient acquisition. Abiotic factors, such as warming and drought, have been shown to influence exudate profiles by modulating enzyme activity and metabolite transport (Yang et al., 2025). Collectively, these findings point to the integrative nature of plant metabolic responses to environmental and microbial contexts.

### Environmental conditions influence fungal diversity, composition, and abundance

Fungal taxa exhibit strong ecological differentiation, often shaped by subtle biotic and abiotic variations that reflect their functional roles and life history traits (Bahram & Netherway, 2022; Bui et al., 2020). These patterns intensify under environmental disturbance, where altered soil chemistry and microbial interactions act as filters, favoring competitively dominant or stress-tolerant taxa (Delgado-Baquerizo et al., 2020). In this study, higher soil moisture was associated with greater fungal diversity, likely due to improved conditions for microbial activity and plant–microbe interactions. This aligns with previous findings showing that water limitation can induce persistent shifts in forest microbial communities (Jaeger et al., 2023). Differential abundance analyses further revealed treatment-driven restructuring of fungal assemblages, consistent with taxon-specific ecological strategies. For instance, Hydnodontaceae was enriched in soil fungicide treatments (likely reflecting reduced AMF), but depleted under foliar fungicide and heat sterilization. This suggests potential dependency on complex microbial networks involving both AMF and pathogens. In contrast, Glomerellaceae increased in heat-treated soils, possibly exploiting reduced microbial competition, but declined under fungicide applications, indicating sensitivity to broader microbial disruptions. These shifts reflect differential responses to altered resource and interaction landscapes. Nectriaceae, a predominantly pathogenic family (Lombard et al., 2015), also responded strongly to biotic and abiotic treatments, with elevated prevalence under altered conditions. This suggests that community restructuring may create ecological opportunities for certain taxa, potentially with implications for plant health and rhizosphere dynamics (Osborne et al., 2018; Taylor et al., 2014). Taken together, these results highlight fine-scale functional differentiation within fungal communities, shaped by variation in stress tolerance, resource use, and microbe–microbe interactions.

### Fungal community structure and metabolite composition are tightly linked

Root exudates shape rhizosphere microbial assembly through the release of primary and secondary metabolites that recruit or deter specific taxa (Leach et al., 2017; Nobori et al., 2018). In this study, *G. guidonia* metabolites were significantly associated with rhizosphere fungal community structure, reinforcing the role of exudates as key mediators of belowground plant–microbe interactions (Broeckling et al., 2008; Zhalnina et al., 2018). Mantel tests revealed significant associations between fungal communities and all major metabolite classes, namely alkaloids, carbohydrates, fatty acids, polyketides, shikimates and phenylpropanoids, amino acids, and terpenoids. This suggests that diverse compound groups contribute to shaping fungal assemblages, supporting the idea that chemically diverse exudates serve as ecological filters that help select for fungi that metabolize or tolerate particular compounds (Clocchiatti et al., 2021; Pang et al., 2021). Positive correlations between metabolite and fungal diversity also suggest that richer chemical environments may support higher microbial diversity, likely through expanded resource niches or signaling pathways (Zhalnina et al., 2018). However, redundancy analysis showed that aggregate metabolomic profiles only explained a small proportion of fungal community variation, implying that specific metabolite–taxon associations rather than global chemical shifts could be the primary ecological drivers. While fungal composition was sensitive to microbe-altering treatments (e.g., fungicides), it was not strongly linked to overall metabolite diversity. This implies an asymmetric relationship where metabolites could be playing a dominant role in structuring early fungal communities although fungal feedbacks may be limited at this seedling stage. Fine-scale pairwise analyses revealed significant associations between individual metabolites and specific fungal OTUs, underscoring the importance of targeted interactions that may be masked in community-level metrics. Taken together, these results point to a close link between rhizosphere fungal communities and metabolite composition. However, given that soil inoculum (microbial legacy) also influenced metabolite profiles, the directionality of these associations is likely context-dependent and may reflect complex feedbacks between plant exudation and microbial community dynamics.

### Metabolite and fungal variation is linked to seedling traits

Although overall treatment effects on seedling traits were modest, we observed significant associations between metabolite clusters, metabolite principal components, and plant morphological traits. Fungal community structure was also significantly correlated with trait variation. These findings suggest that belowground chemical and microbial variation may influence early plant phenotypes, albeit subtly and in a context-dependent manner. Notably, leaf surface area and specific leaf area (SLA) exhibited clear relationships with metabolite composition. Specifically, SLA was positively associated with Cluster 1 metabolites, which was dominated by secondary metabolites, and negatively associated with Clusters 2 and 3 that were primarily enriched in primary metabolites (Fig. S2b). This pattern may reflect shifts in carbon allocation strategy, where seedlings with thinner leaves potentially prioritize the production of secondary metabolites such as phenolics and terpenoids, which have known roles in defense and microbial signaling (Badri & Vivanco, 2009). In contrast, seedlings associated with Clusters 2 and 3 may favor primary metabolism that supports nutrient uptake, microbial facilitation, or root biomass production. These divergent associations suggest possible trade-offs between growth and defense, consistent with broader patterns in plant functional trait ecology (Freschet et al., 2015). Associations with dry root mass followed a similar but less consistent trend, suggesting a potential but weaker role for belowground allocation in shaping exudate profiles. The stronger correlation between metabolite composition and trait dissimilarity, compared to that between metabolite richness and traits, highlights the ecological relevance of qualitative shifts in exudate chemistry. This indicates that variation in the type, rather than the number, of metabolites more strongly influences plant morphological outcomes.

Fungal community composition also correlated with trait dissimilarity, albeit with lower explanatory power, suggesting a more diffuse or indirect role that is potentially mediated by altered microbial signaling or nutrient dynamics. Such effects may reflect indirect or delayed induction of pathways through which rhizosphere composition shapes trait expression, consistent with prior evidence that plant–microbe interactions often intensify over time or under prolonged stress (Ke et al., 2021; Ravanbakhsh et al., 2019). For instance, trait– metabolite divergence was strongest in comparisons involving heat-treated inoculum, indicating a legacy effect of microbial depletion or abiotic stress. Conversely, soil fungicide treatments caused relatively little divergence, suggesting that selective AMF suppression alone had limited influence on trait expression or exudate patterns within the experimental window. Trait–fungal divergence was more broadly distributed but less pronounced, further supporting a primary role for exudate composition in driving morphological outcomes. Overall, our results suggest that seedling trait variation is more closely linked to root exudate composition than rhizosphere fungal diversity, and that treatment history plays a key role in structuring these interactions. Compositional shifts in the exudate profile, particularly those involving secondary metabolites, may represent an underrecognized axis of adaptive plasticity during early plant development. Future work across ontogeny and in field conditions will be essential to resolve the ecological and mechanistic significance of these belowground–aboveground trait linkages.

## Conclusion

This study reveals how environmental conditions reshape plant–soil–microbe interactions by altering both root exudate chemistry and fungal community composition in a tropical tree species. We show that treatment effects, particularly warming-induced microbial legacy and soil moisture, modulate root exudate metabolite profiles and that microbial manipulations significantly shift metabolite composition and fungal structure. These shifts suggest that root exudation is governed by a dynamic interplay between abiotic conditions and microbial context, with metabolites in turn acting as ecological filters that shape fungal assemblages. While our data reveal significant associations between metabolite composition and fungal community structure, links to seedling traits were weak at this developmental stage suggesting that belowground feedbacks influencing plant morphology may be subtle during early growth. It is possible that the association could strengthen over time or under cumulative environmental stress. The responses of distinct fungal families, such as Hydnodontaceae and Nectriaceae, further support the idea of functional filtering driven by environmental and microbial cues. Taken together, our results underscore the need to consider metabolite-mediated pathways in understanding plant resilience to climate change. They highlight the importance of multifactorial approaches that integrate abiotic stress, microbial dynamics, and metabolic plasticity. Future work should examine how these interactions evolve over time, how different microbial guilds contribute, and how specific metabolite pathways influence plant fitness and ecosystem recovery under ongoing global change.

## Acknowledgements

This work was funded by NSF grant DEB 2120085, Department of Energy (DE-SC0012000, DE-SC-0011806, 89243018S-SC-000014, 89243018S-SC-000017, DE-SC-0018942, DE-SC0022095, 89243021S-SC-000076) and NSF (DEB-1754713). We would like to acknowledge the Pennsylvania State University’s Huck Metabolomics Core Facility (RRID:SCR_023864) for use of the Vanquish Horizon UHPLC System and Ashley Shay for helpful discussions on sample preparation. We also thank Shelby McMahan, William Mejía, Alberto Ibáñez, Deyaneira Iglesias, and Laura Rubio for assistance with shade house experiments.

## Author contributions

**JRL**, and **BB** conceptualized and designed the study and acquired funding. **JM**, **PB** and **GHV** performed shade house experiments, with input from **IFG**. **JM** processed samples for metabolomics analysis. **BB** oversaw sample processing for fungal DNA sequencing. **JM** analyzed the data with input from **JRL** and **BB**. **JM** wrote the manuscript. All authors provided feedback on the manuscript.

## Data availability

All data supporting the findings of this study are included in the manuscript and supplementary information files. Code used for data analysis is found at https://github.com/Jmasanga/Guarea_guidonia_analysis

