## Supplemental Figures 1-7 for "Legacy of warming and microbial treatments shape root exudates and rhizosphere fungal communities of a tropical tree"

### **Supplementary figures**

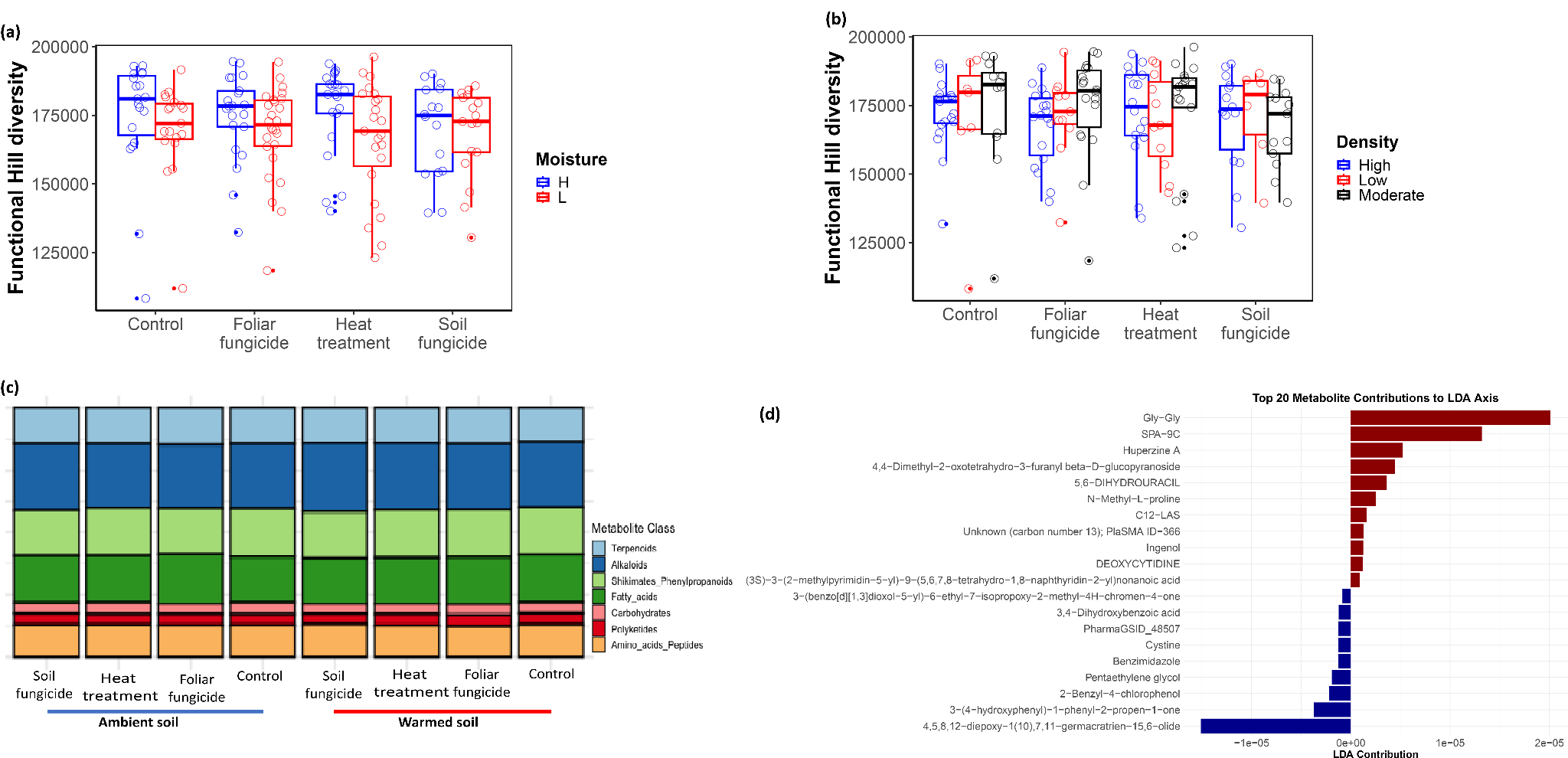

**Fig. S1.**

Root exudate metabolite responses to antimicrobial and abiotic treatments. (a) Functional Hill diversity of metabolites across antimicrobial treatments (x-axis), colored by soil moisture (High [H] vs. Low [L]). (b) Functional Hill diversity across microbial treatments colored by plant density (High, Moderate, Low). (c) Relative abundance of metabolite classes across antimicrobial treatments and microbial legacy (inoculum source). (d) Linear discriminant analysis (LDA) of metabolite features that distinguish treatments. The top 20 contributors are shown. Red and blue colors denote positive and negative contributions, respectively.

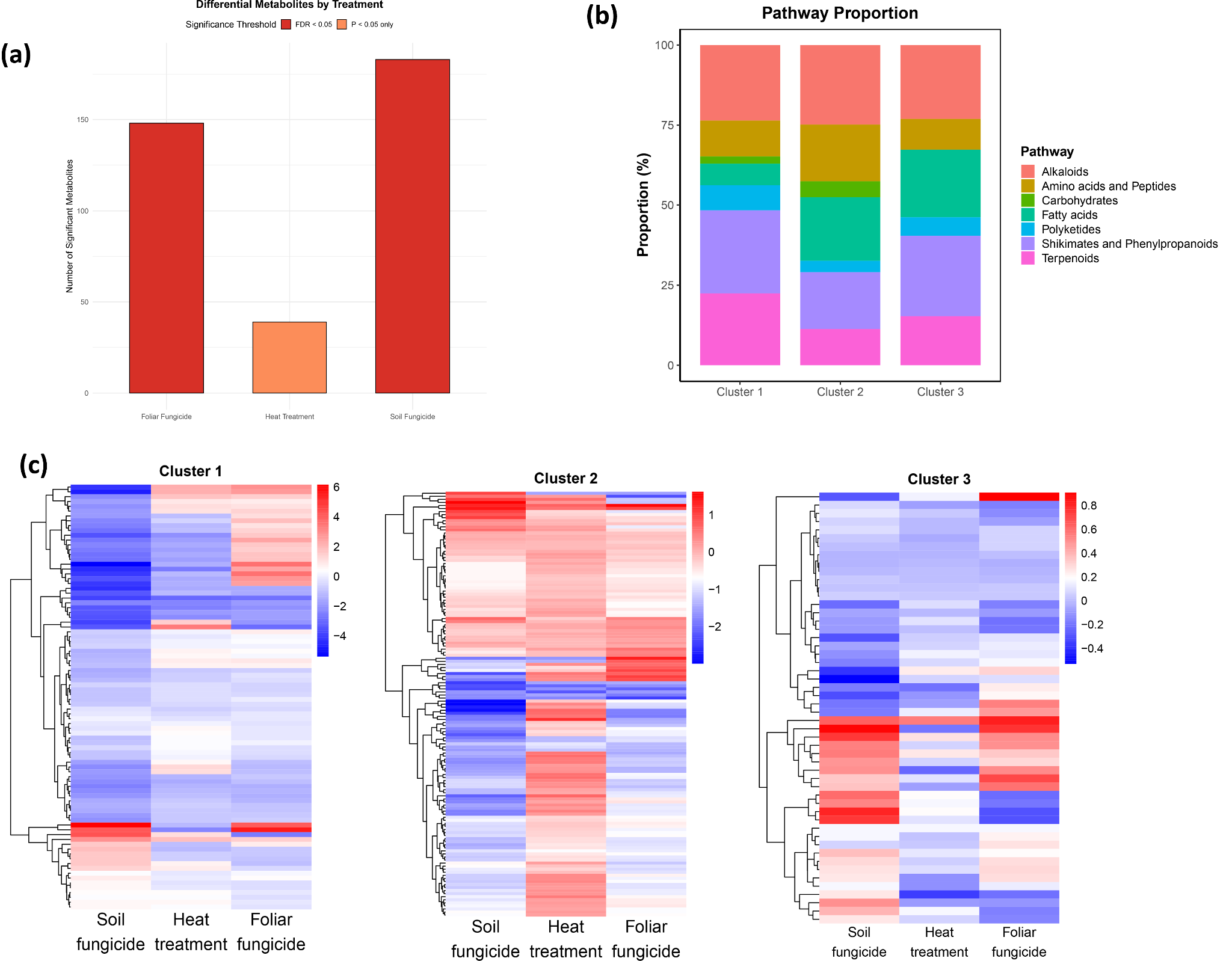

**Fig. S2.**

Treatment-specific modulation of root exudate metabolites. (a) Number of significantly altered metabolite features across microbe-altering treatments, identified using the “limma” package. Bars depict counts of differentially abundant metabolites relative to controls (no treatments to soil inoculum). (b) Proportions of metabolite pathways across three consensus clusters of co-occurring features. (c) Heatmaps of standardized metabolite peak intensities (Z-score) for the three consensus clusters across antimicrobial treatments.

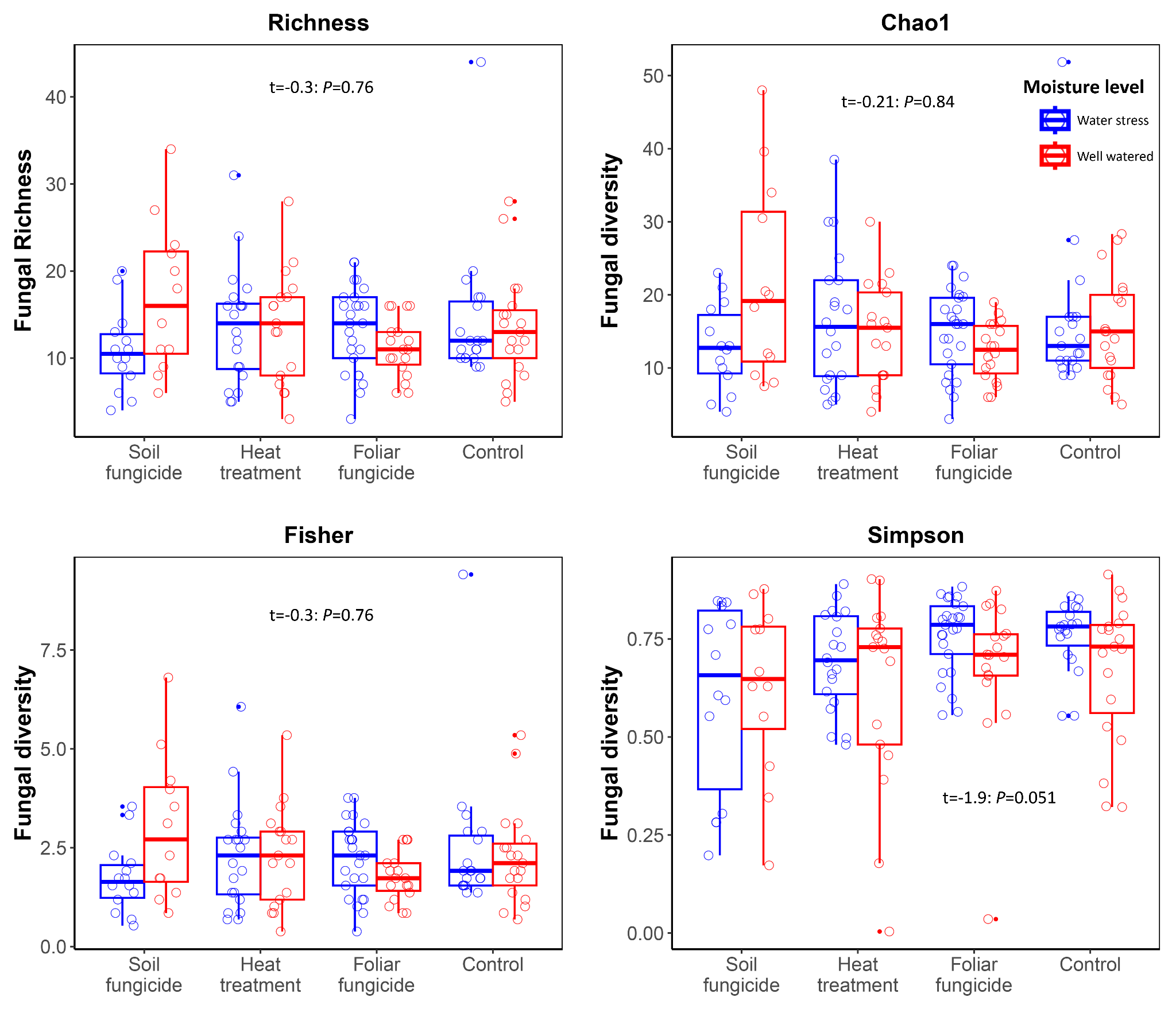

**Fig. S3.**

Fungal alpha diversity across antimicrobial treatments and moisture levels.

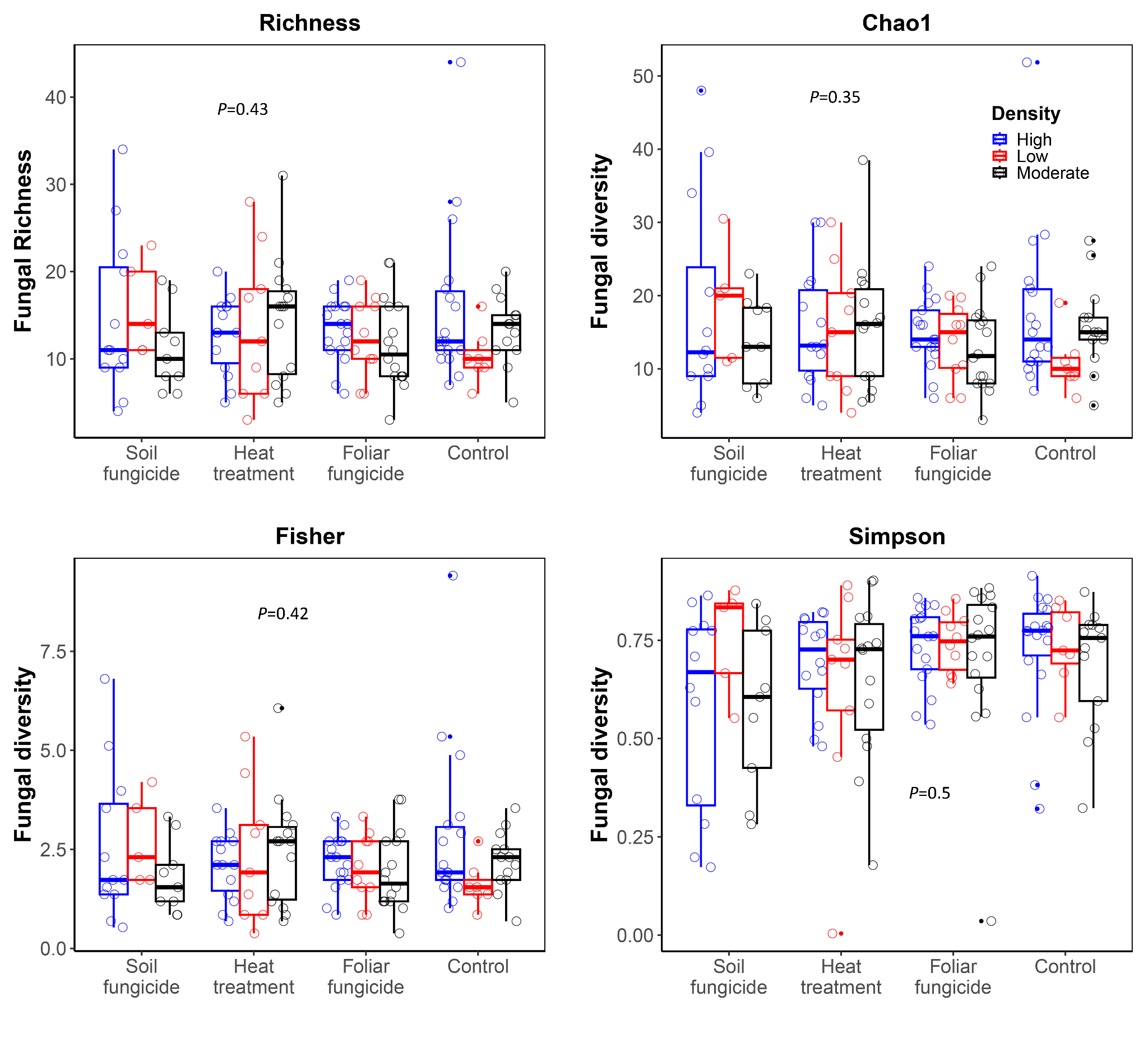

**Fig. S4.**

Fungal alpha diversity in the rhizosphere of *G. guidonia* across antimicrobial treatments and plant density. Low, moderate, and high densities represent 1, 3 and 5 plants per pot, respectively.

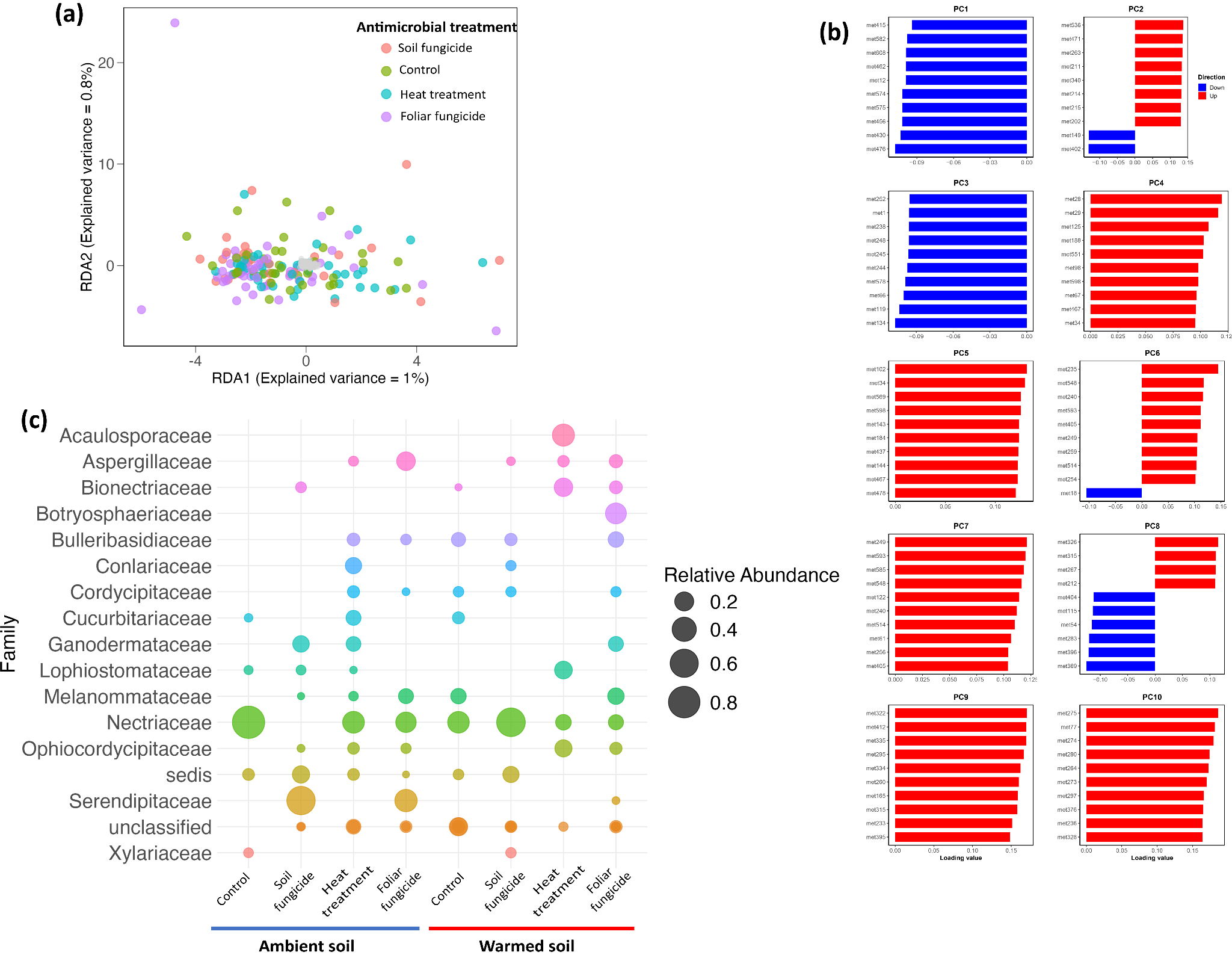

**Fig. S5.**

Associations between exudate metabolites and rhizosphere fungal communities. (a) RDA of fungal community composition constrained by root exudate metabolite profiles across 144 paired samples (individual points colored by antimicrobial treatment). (b) Loadings for the top 10 metabolite PCs contributing to the constrained axes. (c) Bubble plot showing relative abundance of outlier fungal families compared across antimicrobial treatments. Outlier taxa were identified based on Mahalanobis distance from the multivariate centroid in RDA space. Bubble sizes denote the degree of multivariate deviation from the centroid.

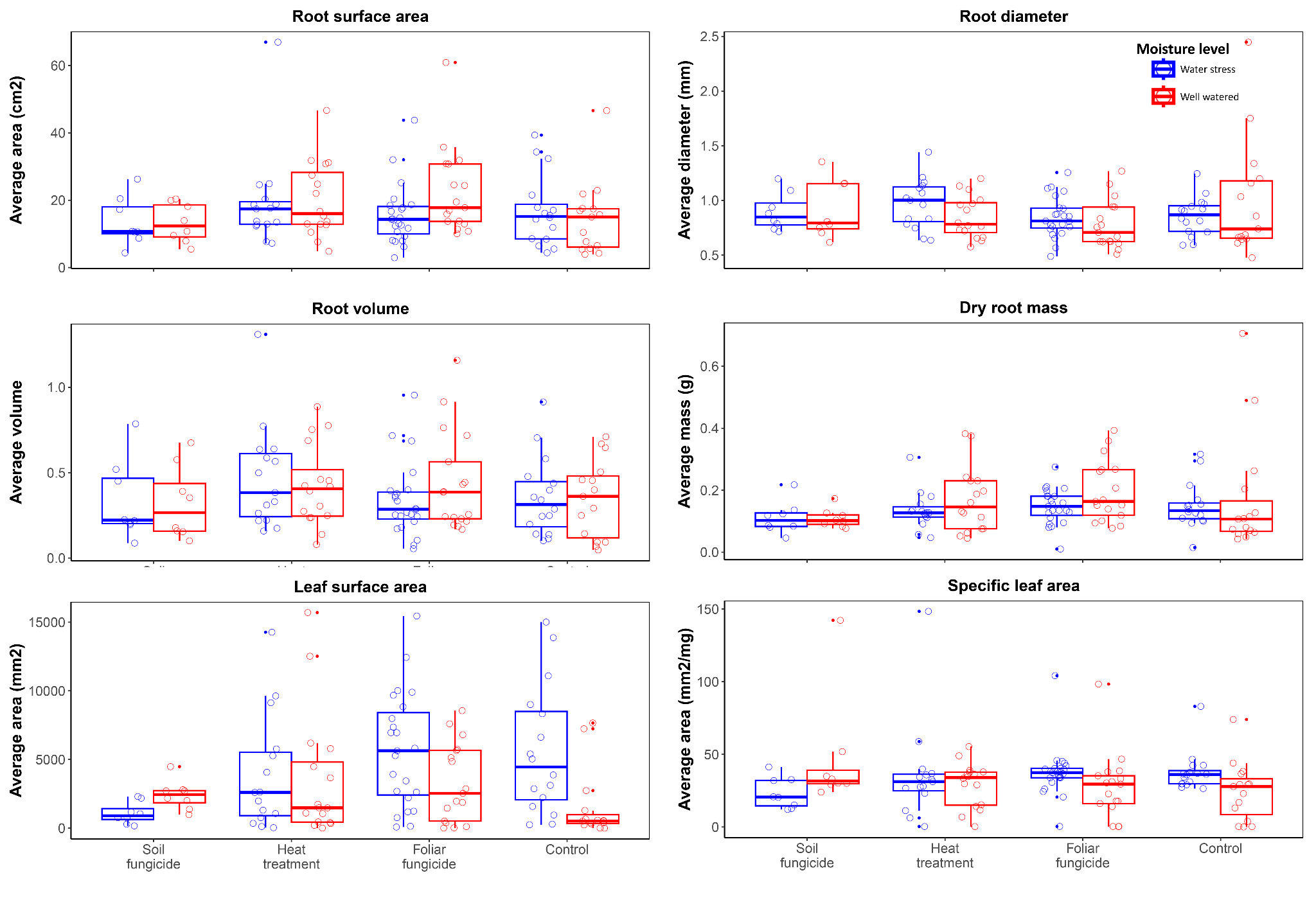

**Fig. S6.**

Seedling trait responses to microbial-altering treatments (x-axis) and colored by water availability. Points in the boxplots represent biological replicates. All traits, with the exception of dry root mass, showed no significant response to treatments (p > 0.05). Statistical significance was assessed using linear models accounting for all treatment factors.

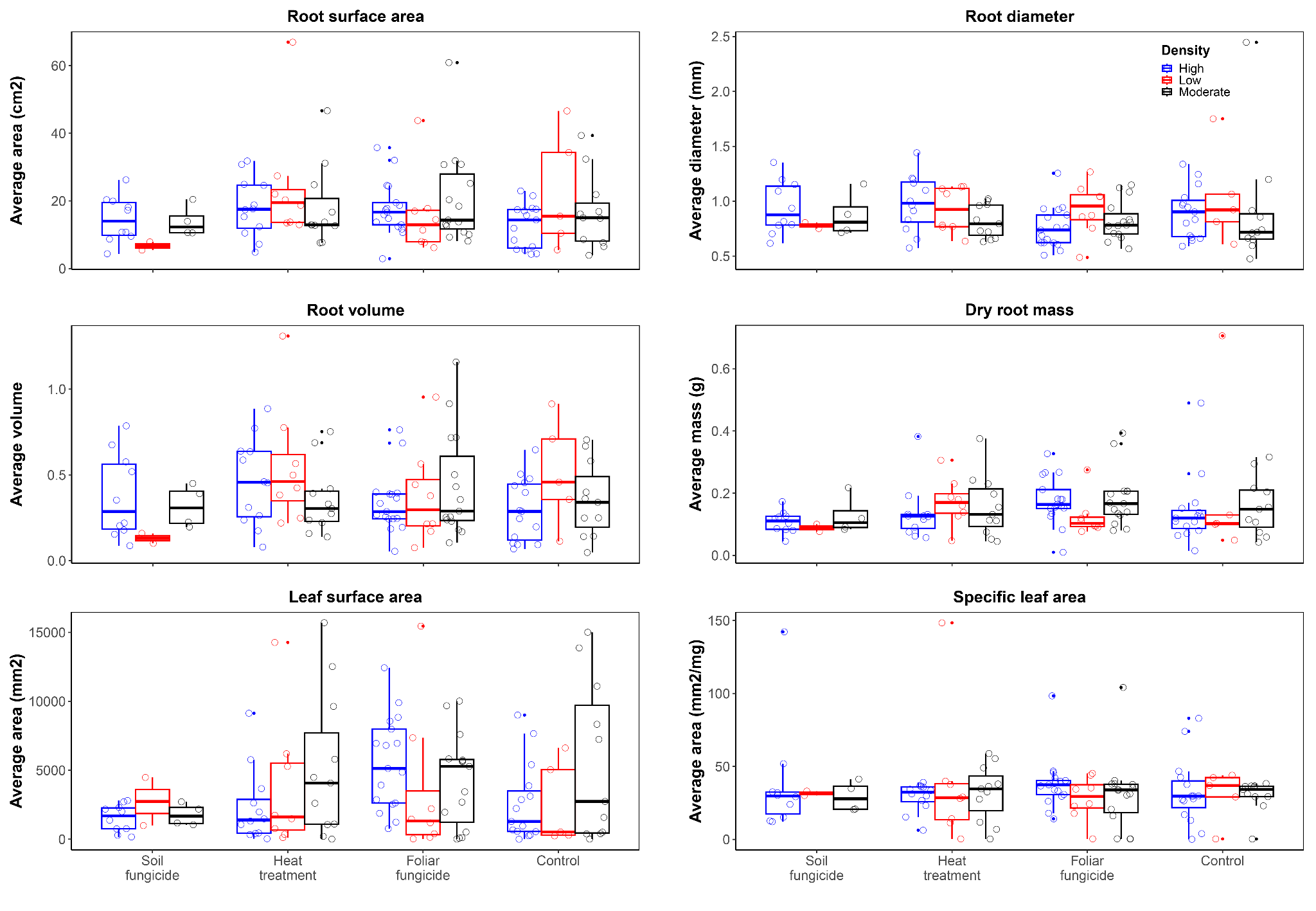

**Fig. S7.**

Seedling trait responses to microbial-altering treatments (x-axis) and plant density (color). Density is defined as low (1 plant per pot), moderate (3 plants), and high (5 plants). Points are biological replicates. All traits, apart from dry root mass, showed no significant response to treatments (p > 0.05). Statistical significance was assessed using linear models accounting for all treatment factors.
