## Supplemental Tables 1 & 2 for "Legacy of warming and microbial treatments shape root exudates and rhizosphere fungal communities of a tropical tree"

### **Supplementary tables**

**Table S1.** Metabolite cluster associations with fungal diversity and seedling traits.

| **Cluster** | **Variable** | **rho** | **Pvalue (raw)** | **Pvalue (adjusted)** |
| --- | --- | --- | --- | --- |
| Cluster1 | Shannon | -0.1039 | 0.2609 | 0.2609 |
| Cluster2 | Shannon | 0.2483 | 0.0065 | **0.0194** |
| Cluster3 | Shannon | 0.2316 | 0.0113 | **0.0225** |
| Cluster1 | LSA | -0.3254 | 0.0003 | **0.0009** |
| Cluster2 | LSA | 0.2767 | 0.0023 | **0.0046** |
| Cluster3 | LSA | 0.2623 | 0.0040 | **0.0046** |
| Cluster1 | SLA | -0.1981 | 0.0308 | 0.0923 |
| Cluster2 | SLA | 0.1409 | 0.1263 | 0.1796 |
| Cluster3 | SLA | 0.1562 | 0.0898 | 0.1796 |
| Cluster1 | RSA | -0.1426 | 0.1220 | 0.3659 |
| Cluster2 | RSA | 0.0012 | 0.9897 | 0.9897 |
| Cluster3 | RSA | 0.1014 | 0.2724 | 0.5449 |
| Cluster1 | DRM | -0.1852 | 0.0438 | 0.1313 |
| Cluster2 | DRM | 0.1490 | 0.1057 | 0.1313 |
| Cluster3 | DRM | 0.1738 | 0.0588 | 0.1313 |
| Cluster1 | Root volume | -0.0852 | 0.3568 | 1.0000 |
| Cluster2 | Root volume | -0.0718 | 0.4377 | 1.0000 |
| Cluster3 | Root volume | 0.0554 | 0.5497 | 1.0000 |
| Cluster1 | ARD | -0.0124 | 0.8937 | 1.0000 |
| Cluster2 | ARD | -0.0993 | 0.2825 | 0.8474 |
| Cluster3 | ARD | 0.0072 | 0.9384 | 1.0000 |

Correlations are between metabolite consensus cluster values with fungal Shannon diversity and seedling morphological traits, performed using Spearman’s rank test. Bold numbers indicate significant associations after multiple testing (adjusted P< 0.05) based on false discovery rate (FDR). LSA- leaf surface area; SLA- specific leaf area; RSA- root surface area; DRM- dry root mass; ARD- average root diameter.

**Table S2:** Effects of experimental treatments on seedling morphological traits

| **Trait** | **F-statistic** | **P-value** |
| --- | --- | --- |
| Specific leaf area | 1.497 | 0.07 |
| Leaf surface area | 1.282 | 0.17 |
| Dry root mass | 2.707 | **8.93e-05** |
| Root surface area | 1.165 | 0.28 |
| Root diameter | 1.041 | 0.43 |
| Root volume | 0.8146 | 0.76 |

Linear models were used to test effects of origin of soil inoculum, antimicrobial treatments, soil moisture, and plant density. The significant effect (p < 0.05) is bolded.
